# Integrated in vivo quantitative proteomics and nutrient tracing reveals age-related metabolic rewiring of pancreatic β-cell function

**DOI:** 10.1101/451542

**Authors:** Matthew Wortham, Jacqueline R. Benthuysen, Martina Wallace, Jeffrey N. Savas, Francesca Mulas, Ajit S. Divakaruni, Fenfen Liu, Verena Albert, Brandon L. Taylor, Yinghui Sui, Enrique Saez, Anne N. Murphy, John R. Yates, Christian M. Metallo, Maike Sander

**Author notes:** These authors contributed equally (Matthew Wortham, Jacqueline R. Benthuysen).

## Abstract

Pancreatic β-cell physiology changes substantially throughout life; yet, the mechanisms that drive these changes are poorly understood. Here, we performed comprehensive in vivo quantitative proteomic profiling of pancreatic islets from adolescent and one-year-old mice. The analysis revealed striking differences in abundance of enzymes controlling glucose metabolism. We show that these changes in protein abundance are associated with higher activities of glucose metabolic enzymes involved in coupling factor generation as well as increased activity of the coupling factor-dependent amplifying pathway of insulin secretion. Nutrient tracing and targeted metabolomics demonstrated accelerated accumulation of glucose-derived metabolites and coupling factors in islets from one-year-old mice, indicating that age-related changes in glucose metabolism contribute to improved glucose-stimulated insulin secretion with age. Together, our study provides the first in-depth characterization of age-related changes in the islet proteome and establishes metabolic rewiring as an important mechanism for age-associated changes in β-cell function.

## Introduction

It has long been recognized that islet β -cells undergo changes in glucose-stimulated insulin secretion (GSIS) with age. Recent studies of rodent and human islets have shown an age-dependent increase in GSIS when juvenile islets are compared to islets during middle age or later in life (Arda et al., 2016; Avrahami et al., 2015; Gregg et al., 2016). Consistent with the observed islet-intrinsic changes to GSIS, circulating insulin levels in both the fasted state and in response to a glucose challenge are higher in older animals (Avrahami et al., 2015; Gregg et al., 2016). These age-dependent functional changes may reflect both maturation and aging processes, defined as those preceding or following sexual maturity, respectively. Several mechanisms have been proposed to be responsible for the increase in GSIS with age, including increased expression of transcription factors that regulate insulin secretory genes in β -cells (Arda et al., 2016; Avrahami et al., 2015) as well as activation of a senescence program by the cell cycle inhibitor p16^Ink4a^ (Helman et al., 2016). However, our current knowledge of age-associated changes in β -cells is largely based on transcriptome studies and an understanding of how age affects the abundance of proteins is lacking.

Studies of the islet proteome could provide mechanistic insights into how age affects islet cell function, but these studies have been technically challenging due to the need for large protein amounts and the limited islet material that can be isolated from rodents. A further challenge of proteomic experiments is comprehensive coverage of the cell proteome, because liquid chromatography-tandem mass spectrometry (LC-MS/MS) systems used in proteomics tend to bias detection toward the most abundant proteins. Recent advances in proteomics combining stable isotope labeling of amino acids in mammals (SILAM) with multidimensional protein detection technology (MudPIT) (McClatchy et al., 2007; Washburn et al., 2001) have helped overcome these obstacles. This approach has recently provided mechanistic insight into the long-lived proteins of the aging brain (Savas et al., 2012). To date, age-related changes in the islet proteome have not been studied. Thus, the impact of transcriptional changes on protein abundance has yet to be broadly determined, and the contribution of posttranscriptional regulation to age-associated functional changes of pancreatic islets remains to be characterized.

Insulin secretion is intimately linked to the rate of β -cell glucose metabolism. Therefore, determining how β -cell glucose metabolism changes throughout life could provide important insight into the mechanisms that mediate the age-associated change in GSIS. The workhorse model for metabolomic studies of β -cells has been the INS1 832/13 insulinoma cell line (Alves et al., 2015; Lorenz et al., 2013; Mugabo et al., 2017). Glucose tracing experiments and measurements of metabolite abundance during glucose stimulation have helped identify β -cell-characteristic patterns of glucose utilization as well as candidate metabolites involved in the regulation of GSIS (Alves et al., 2015; Farfari et al., 2000; Lorenz et al., 2013; Lu et al., 2002; Mugabo et al., 2017). Nutrient tracing has also been employed in primary islets to monitor specific metabolic reactions (Adam et al., 2017; Li et al., 2008; Wall et al., 2015). However, islet nutrient metabolism has not been broadly characterized and it is unknown whether β -cells regulate GSIS with age by altering glucose metabolism.

In this study, we employed in vivo SILAM MS to comprehensively assess differences in islet protein levels between juvenile and adult mice. We further employed targeted metabolomics coupled with nutrient tracing in islets from juvenile and adult mice and characterized metabolic processes contributing to insulin secretion. Combined, these studies revealed hitherto unknown changes in the abundance of metabolic enzymes with age, which coincide with increased generation of glucose-derived coupling factors involved in the regulation of GSIS. Together, this study provides the first in-depth characterization of age-dependent changes in the islet proteome and establishes metabolic rewiring as an important mechanism for regulating GSIS throughout life.

## Results

### Protein quantification in islets from juvenile and adult mice by in vivo proteomics

To identify age-regulated proteins in pancreatic islets, we performed ^15^N SILAM MudPIT LC-MS/MS (McClatchy et al., 2007; Washburn et al., 2001). We metabolically labeled C57BL/6 mice by administering chow containing ^15^N-labeled amino acids to 3-week-old mice immediately after weaning for 10-11 weeks. Islets were then isolated from these mice (n=56), pooled, and total protein lysates were mixed 1:1 with protein from ^14^N-non-labeled islets from either 4-week-old (n=62) or 1-year-old mice (n=38) (**Figure 1A**), hereafter referred to as “juvenile” mice corresponding to the pre-pubescent stage and “adult” mice corresponding to middle age, respectively. We chose these ages to correspond to time points used in other studies assessing age-dependent changes to the β-cell epigenome and transcriptome (Avrahami et al., 2015) coincident with well-established differences in GSIS (Helman et al., 2016). Proteins were proteolytically digested, loaded onto a two-dimensional column, and analyzed by multidimensional LC coupled to an electro-spray ionization tandem mass spectrometer. The mass spectrometer distinguished “heavy” ^15^N-labeled peptides from “light” ^14^N-non-labeled peptides and, through subsequent protein identification and ratio-of-ratios analyses, we quantified relative protein abundance, comparing juvenile and adult islets (Park et al., 2006).

**Figure 1.**
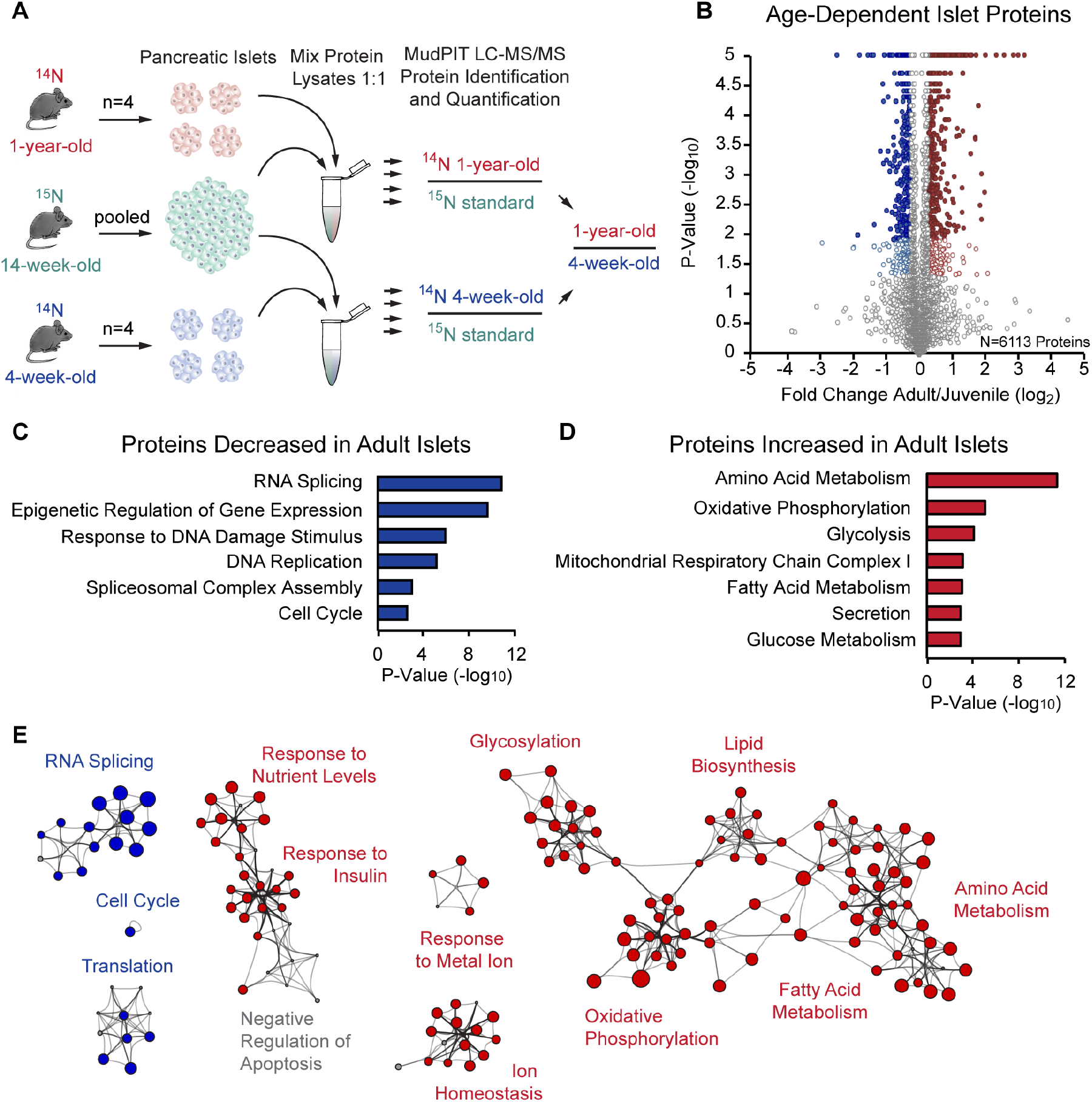
SILAM MudPIT MS comparing protein abundance in juvenile and adult islets. (A) Experimental overview for quantitative analysis of the islet proteome in 4-week-old and 1-year-old mice (n=4 biological replicates per group). (B) Volcano plot comparing protein abundance in juvenile and adult islets; x axis = log2 fold change (adult/juvenile), y axis = -log10 p-value, N = 6,113 total proteins represented: 340 (5.6%) regulated proteins (p-value < 0.05 by t-test and ≥ 1.2-fold change; blue and (A) red open circles) and 1068 (17.5%) most-confident proteins (Benjamini-Hochberg corrected p-value < 0.05 and ≥ 1.2-fold change; blue and red filled circles). p-values = 0 were graphed as 0.00001. Blue circles = downregulated with age, red circles = upregulated with age, grey circles = not regulated with age. (C and D) Gene Ontology (GO) analysis of islet proteins differentially expressed with age. Blue bars = proteins decreased with age, red bars = proteins increased with age. (E) Network of age-regulated islet proteins shown as clustered functional categories. Protein associations were determined using STRING and each node represents a functional category enriched in the protein data as scored by Metascape. Size of node represents extent of enrichment in juvenile or adult. Blue circles = protein categories decreased with age, red circles = protein categories increased with age, grey circles = protein categories that have equal number of proteins up and down with age. See also Figure S1 and Tables S1 and S2.

We achieved ^15^N enrichment of greater than 95.6% (**Figure S1A**). We identified 37,721 peptides (9058 proteins) in 4-week-old islets and 38,657 peptides (9000 proteins) in 1-year-old islets at a false discovery rate (FDR) <1% based on the target-decoy strategy, and quantified 10,245 proteins (**Table S1A**, see Methods). Gene Ontology (GO) analysis of all quantified proteins revealed a broad representation of proteins from multiple cellular components and association with diverse cellular functions and biological processes in a distribution similar to that of all genes (**Figures S1B and S1C**). This demonstrates that our quantitative proteomics approach captured a comprehensive, unbiased, and diverse set of islet proteins.

To identify statistically significant differentially expressed proteins, we generated ^14^N-4-week-old/^15^N-14-week-old or ^14^N-1-year-old/^15^N-14-week-old peptide ratios to calculate a ^14^N-1-year-old/^14^N-4-week-old “ratio of ratios” (**Figure 1A**). We compared four biological replicates of juvenile and adult islets to calculate log2(adult/juvenile) and t-test p-values (**Figure 1B**). Consolidating isoforms, 393 uniquely named proteins were significantly enriched in 4-week-old islets and 551 proteins in 1-year-old islets, exhibiting at least a 1.2-fold change (**Tables S1B and S1C**).

Among the proteins enriched in islets from adult mice was Urocortin 3 (Ucn3) (**Table S1C**), a previously identified β-cell maturation marker involved in the regulation of insulin secretion (Blum et al., 2012; van der Meulen et al., 2015). Thus, our proteomics approach identified changes in protein abundance that are consistent with previous studies.

### Proteins involved in nutrient metabolism are age-dependently regulated in islets

To understand the dynamics of age-dependent regulation of the islet proteome, we performed Gene Ontology (GO) annotation and Gene Set Enrichment Analysis (GSEA) of the differentially expressed proteins. Additionally, we built a network of functional categories by linking proteins via STRING interactions and assigning pathway categories to groups of proteins using Metascape (see Methods) (Szklarczyk et al., 2015; Tripathi et al., 2015). Proteins that decreased in abundance with age were functionally associated with RNA splicing, epigenetic regulation of gene expression, response to DNA damage, translation, and cell cycle regulation (**Figures 1C, 1E and S1D; Tables S2A and S2C**), which is consistent with declining β-cell proliferation during this time period (Teta et al., 2005). Conversely, there was a striking age-dependent increase in proteins associated with the regulation of aspects of cell metabolism, including amino acid metabolism, oxidative phosphorylation, glycolysis, and fatty acid metabolism (**Figures 1D, 1E and S1E; Tables S2B and S2D**). Among the regulators of nutrient metabolism that increased in abundance with age were the enzymes involved in glucose metabolism Aldoa and Eno2, components of the TCA cycle (i.e. Pdha1, Cs, Fh1) and respiratory chain (i.e. Ndufv3, Cox4i1, Atp6v1h), as well as enzymes important for fatty acid (i.e. Acaa2, Acat1, Hadh) and amino acid metabolism (i.e. Bcat2, Bckdhb) (**Table S2B**). Given the coupling of insulin secretion to β-cell nutrient metabolism (Prentki et al., 2013), this suggests that the observed increase in β-cell GSIS with age (Avrahami et al., 2015; Gregg et al., 2016; Helman et al., 2016) could be caused by age-related changes in nutrient metabolism.

### Posttranscriptional regulation accounts for the majority of age-associated changes in islet protein levels

To determine the extent of correlation between the age-regulated islet proteome and transcriptome, we performed RNA-seq on islets from 4-week-old and 1-year-old mice. We found 1348 genes that increased and 1542 genes that decreased in expression with age (≥1.2-fold change, p<0.05; Figure 2A; Table S3). GO and GSEA analysis showed some, but incomplete, overlap between functional categories regulated at the mRNA and protein level (**Figures S2A-D; Table S4**). To assess to which extent individual mRNAs and proteins are co-regulated with age, we calculated the correlation coefficient between changes in protein and mRNA levels. Considering all 10,245 proteins quantified by in vivo proteomics (**Table S1A**), we found a modest correlation between age-associated regulation at the mRNA and protein level (ρ=0.4, p=2.2×10^−16^; Figure 2B). Of the unique proteins that changed in abundance with age, only 11.5% (109 out of 944) were also regulated at the mRNA level (**Figures S2E and S2F; Table S5**). GSEA of differentially-expressed proteins not regulated at the mRNA level revealed involvement in RNA splicing, cell cycle, and proteasome regulation for proteins decreasing with age and oxidative phosphorylation, TCA cycle, and fatty acid metabolism for proteins increasing with age (**Figure 2C; Table S6**). To further validate these findings, we queried our islet proteome network against corresponding mRNA changes by building a similar mRNA category network and identifying the category nodes that overlap with the protein network (**Figure 2D**). This revealed that many categories, including RNA splicing, cell cycle, translation, response to insulin, and ion homeostasis, were mostly exclusive to the protein network (**Figure 2E**). Furthermore, several categories such as oxidative phosphorylation, glycosylation and response to nutrient levels were enriched in the protein network with very few nodes enriched in the mRNA network. In sum, this analysis shows significant age-related changes in the abundance of proteins involved in nutrient metabolism not associated with corresponding transcriptional changes.

**Figure 2.**
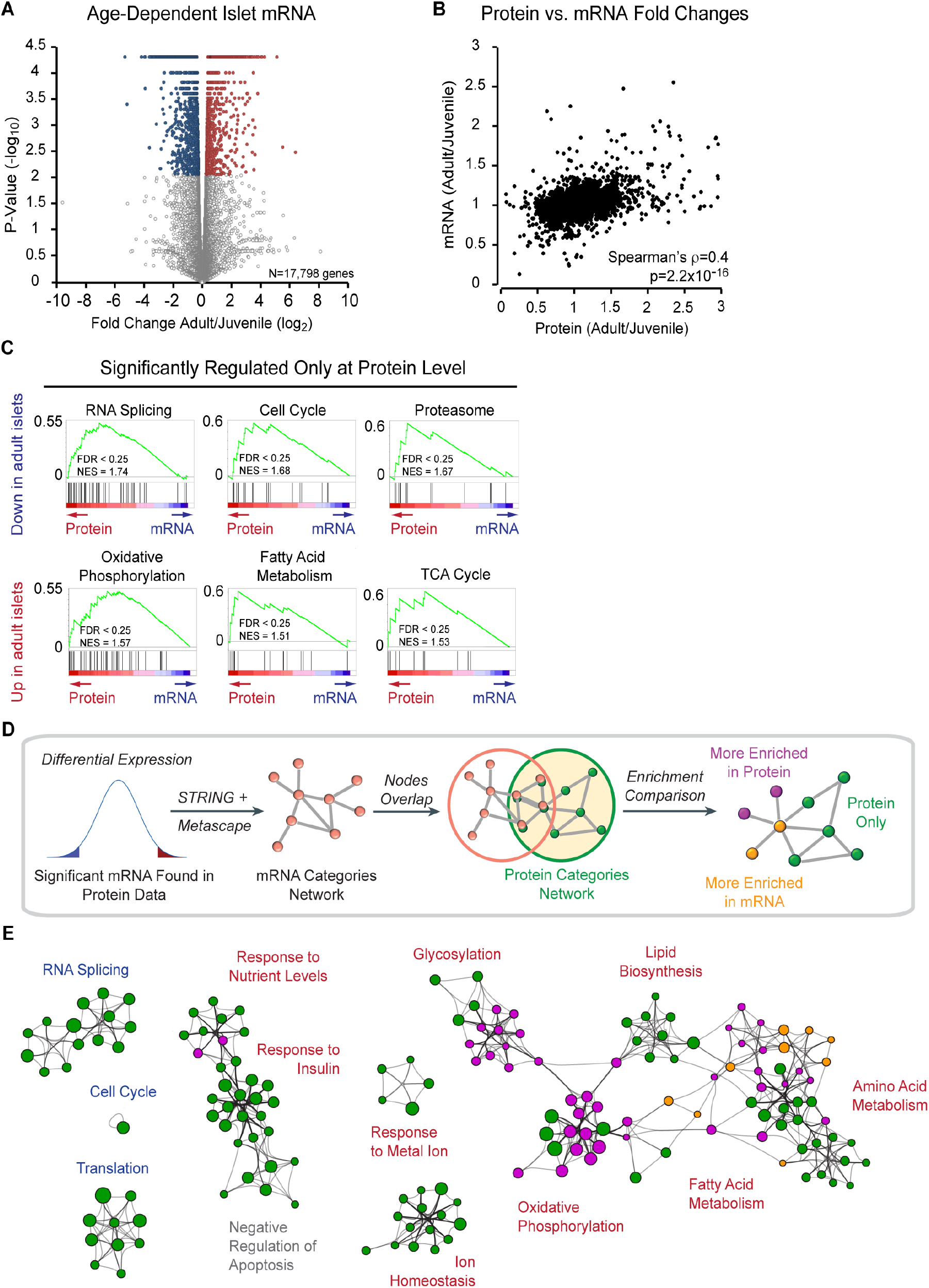
Age-dependent changes in the islet proteome independent of transcriptional regulation. (A) Volcano plot comparing mRNA abundance by RNA-seq in islets from 4-week-old and 1-year-old mice (n=3 biological replicates per group); x axis = log2 fold change (adult/juvenile), y axis = -log10 p-value, N = 17,798 total genes represented. 2896 (16.3%) regulated mRNAs (p-value < 0.05 by unpaired two tailed t-test and ≥ 1.2-fold change; blue circles = downregulated with age, red circles = upregulated with age, grey open circles = not regulated with age). (B) Spearman’s rank correlation between mRNA and protein ratios from juvenile and adult islets (ρ=0.4, p=2.2×10^−16^). (C) Gene set enrichment analysis (GSEA) comparing proteins that change with islet age to corresponding mRNAs. Gene sets predominantly regulated at the protein level show a left-skewed Enrichment Score distribution. (D) Workflow for comparing protein and mRNA networks. (E) Network of clustered functional categories showing protein and mRNA network comparison. Protein associations were determined using STRING and each node represents a functional category enriched in the protein data as scored by Metascape. Size of node represents extent of enrichment in protein or mRNA network, i.e. larger purple circles are categories enriched in protein network to a greater extent than in the mRNA network. Green circles = functional categories only found in protein network, purple circles = functional categories more enriched in the protein network, yellow circles = functional categories more enriched in mRNA network. Blue text = protein categories decreased with age, red text = protein categories increased with age, grey text = protein categories with same enrichment significance in the protein and mRNA data. See also Figure S2 and Tables S3-S6.

### Activity of the amplifying pathway of GSIS increases with age in mice

Insulin secretion is modulated by diverse metabolic inputs (Prentki et al., 2013). While glucose is the key stimulus for insulin secretion, the amplitude of the GSIS response is significantly augmented by metabolites derived from lipid catabolism and the TCA cycle. Having observed that TCA cycle and fatty acid metabolism enzymes are more abundant in older mice, we predicted that these metabolic pathways could contribute to the observed increase in GSIS in aged islets (Avrahami et al., 2015; Gregg et al., 2016; Helman et al., 2016). Therefore, we measured insulin content and secretion in islets from 4-week-old and 1-year-old mice co-stimulated with glucose and nutrients contributing to lipid metabolism (palmitic acid) or the TCA cycle (leucine and glutamine). Total insulin content did not differ between islets from juvenile and adult mice (**Figure S3A**). Consistent with prior findings (Avrahami et al., 2015; Gregg et al., 2016; Helman et al., 2016), islets from 1-year-old mice exhibited a more robust insulin secretory response to high glucose than islets from 4-week-old mice (**Figure 3A**). Co-stimulation with glucose and the amino acids leucine and glutamine potentiated GSIS, but the response was modestly reduced rather than increased in adult compared to juvenile islets (**Figure 3A**; 1.5-fold increase in adult versus 2.2-fold increase in juvenile compared to glucose alone). Potentiation of GSIS by the fatty acid palmitate was absent in islets from 1-year-old mice (0.8-fold in adult versus 2.6-fold in juvenile) (**Figure 3A**). Therefore, alterations in fatty acid and amino acid metabolism are unlikely to explain age-associated differences in GSIS.

**Figure 3.**
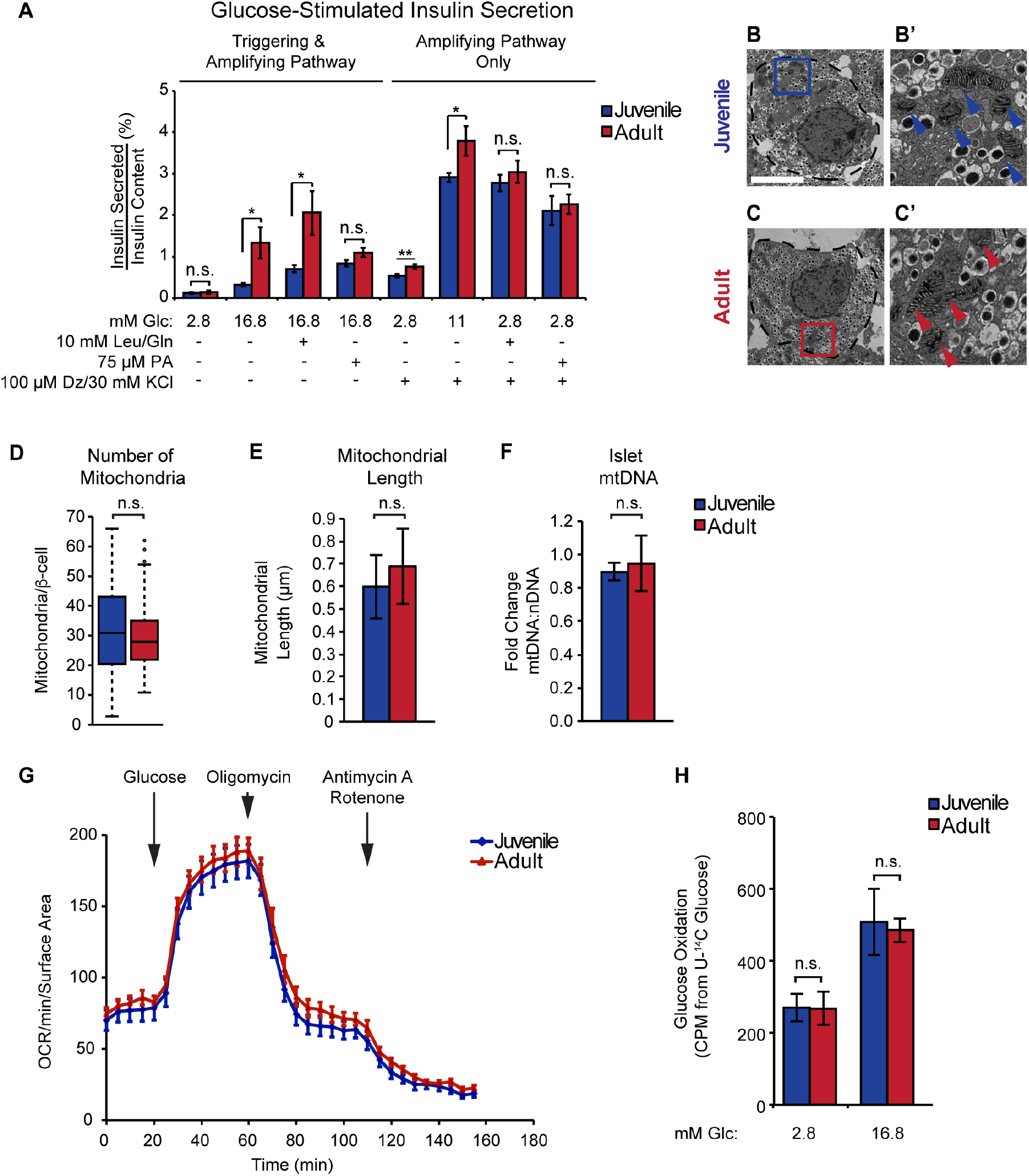
Activity of the amplifying pathway of glucose stimulated insulin secretion increases with age independent of altered respiration. (A) Insulin secretion assay on islets from 4-week-old and 1-year-old mice (n=7-18 per group). Assay was performed in varying glucose (Glc) concentrations as indicated or in conjunction with 10 mM Leucine (Leu)/Glutamine (Gln), 75 µM Palmitate (PA), or 100 µM Diazoxide (Dz) and 30 mM KCl. (B and C) Electron micrograph showing a juvenile and adult β-cell (outlined by dotted line). Boxed areas in B and C are shown in B‘ and C’ at 8x magnification. Arrows indicate mitochondria. Scale bar = 5 µm. (D) Quantification of mitochondria in juvenile and adult β-cells (n=50-60 β-cells from 6 mice). (E) Quantification of mitochondrial length (n=49-59 mitochondria from 10 β-cells). (F) Mitochondrial DNA (mtDNA) compared to nuclear DNA (nDNA) from juvenile and adult islets (n=3 mice per group). (G) Oxygen consumption rates (OCR) of islets from juvenile and adult mice treated with 16.8 mM glucose, 5 µM Oligomycin, 0.5 µM Rotenone and 2 µM Antimycin A (n=7-10 per group). (H) Glucose oxidation rates in juvenile and adult islets as determined by release of 14CO2 from islets incubated with U-14C glucose. Islets were incubated with the indicated concentrations of unlabeled glucose (Glc) spiked with U-14C glucose (n=5-6 per group). Error bars represent ± S.E.M.; *p < 0.05 by unpaired two tailed t-test; n.s., not significant. See also Figure S3.

Nutrients can potentiate insulin secretion through inhibition of the KATP channel by serving as substrates for ATP synthesis (the triggering pathway) or through production of intermediary metabolites that enhance the effect of Ca^2+^ on insulin secretion (the amplifying pathway). Metabolites capable of activating the amplifying pathway are termed metabolic coupling factors. To assess the ability of nutrients to enhance insulin secretion independent of KATP channel closure, we provided islets with glucose, amino acids, or fatty acids, while simultaneously treating with KCl and diazoxide to stimulate Ca^2+^ influx and prevent further changes in membrane potential through the KATP channel (**Figure 3A**). Under these conditions, islets from adult mice secreted more insulin in response to glucose than islets from juvenile mice, whereas the two groups responded similarly to amino acids and palmitic acid. Similar responses of adult and juvenile islets to amino acids and palmitate during forced depolarization (measuring the amplifying pathway alone) are in contrast to age-related differences in the ability of these nutrients to potentiate insulin secretion in response to glucose (which reflects both triggering and amplifying pathways). These observations suggest there could be modest age-dependent differences in modulation of the triggering pathway of insulin secretion by these nutrients. Altogether, these results indicate that enhanced insulin secretion of adult islets reflects a specific hypersensitivity to glucose rather than a heightened response to nutrients in general. Furthermore, the increased glucose responsiveness of adult islets occurs in part through mechanisms independent of KATP channel inhibition, suggesting that changes in glucose metabolism leading to altered production of metabolic coupling factors could underlie enhanced GSIS with age.

The mitochondrion is a major site for the generation of metabolic coupling factors during glucose stimulation (Prentki et al., 2013). Given the upregulation of proteins involved in oxidative phosphorylation with age, we predicted that increased mitochondrial abundance or respiratory capacity could underlie the generation of metabolites that stimulate the amplifying pathway of insulin secretion. Thus, we investigated mitochondrial function and morphology in juvenile and adult islets. Transmission electron microscopy (TEM) of β -cells did not reveal an increase in mitochondrial number or size with age (**Figures 3B-E**), nor was mitochondrial DNA (mtDNA) content increased (**Figure 3F**). This was in contrast to the well-established decline of mtDNA in liver with age (**Figure S3B**) (Bratic and Larsson, 2013). In addition, we measured oxygen consumption in islets of juvenile and adult mice and found no difference in mitochondrial respiration during glucose stimulation or following treatment with the ATP synthase inhibitor oligomycin (**Figure 3G**), indicating that mitochondrial ATP synthesis is unchanged with age. Similarly, oxygen consumption rates after leucine and glutamine stimulation as well as maximal respiration rates after treatment with the mitochondrial uncoupler FCCP showed no difference between juvenile and adult islets (**Figure S3C**). Furthermore, tracer experiments using 5-^3^H labeled glucose and uniformly-^14^C labeled glucose (U-^14^C glucose) revealed identical rates of glucose utilization and oxidation, respectively, in juvenile and adult islets stimulated with glucose (**Figures 3H and S3D**). Thus, despite increased abundance of mitochondrial proteins in adult islets, mitochondrial respiration was not altered, suggesting that other aspects of the stimulus secretion pathway are contributing to the increase in GSIS with age.

### Tracing the metabolic fate of glucose in primary islets

As an alternative hypothesis, we considered that the generation of coupling factors produced by glycolysis and the TCA cycle under high glucose could be enhanced in adult β-cells. Since studies examining nutrient metabolism in β-cells have mostly been conducted in insulinoma cell lines (Alves et al., 2015; Lorenz et al., 2013; Mugabo et al., 2017), it was first necessary to establish techniques for glucose tracing and targeted metabolomics within intact mouse islets. Our approach aimed to characterize glucose metabolism through glycolysis and the TCA cycle during acute glucose stimulation. To equilibrate metabolism to a basal state, we pre-incubated islets in 2.8 mM unlabeled glucose for one hour, then transferred islets to uniformly-^13^C-labeled glucose (U-^13^C glucose hereafter) at a concentration of 2.8 mM or 16.8 mM glucose for one hour, and quantified the abundance and extent of ^13^C labeling of metabolites by GC-MS (**Figure 4A; Table S7**).

**Figure 4.**
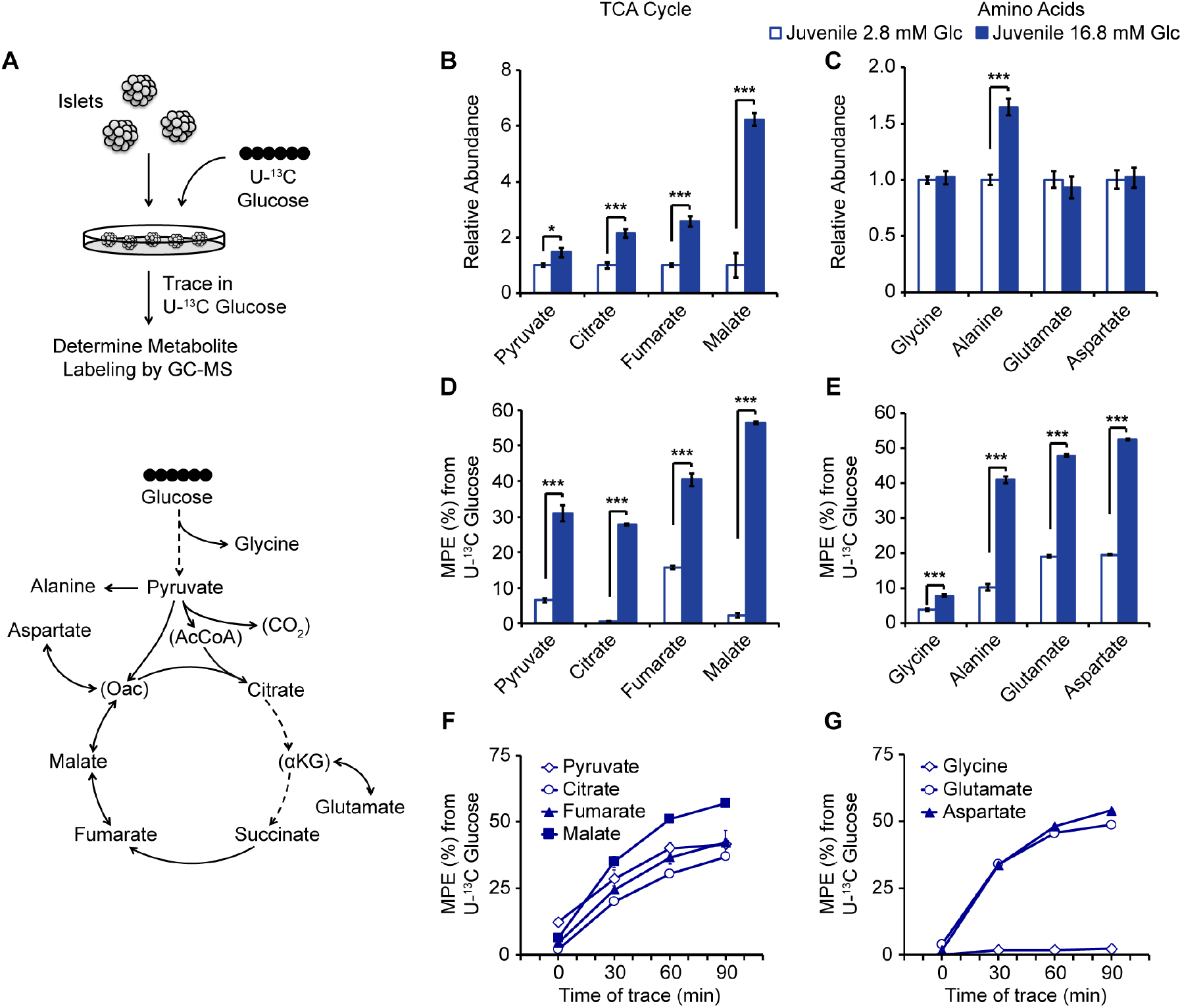
U-^13^C glucose tracing and targeted metabolomics in islets from juvenile mice. (A) Schematic of tracing strategy (top). Overview of intermediary metabolites detected by GC-MS in islets (bottom). Central carbon metabolites below the GC-MS detection limit are indicated in parentheses. (B and C) Abundances of the indicated metabolites following a one-hour trace with 2.8 mM or 16.8 mM U-^13^C glucose in juvenile islets, expressed relative to metabolite levels of islets incubated in 2.8 mM U-^13^C glucose. (D and E) Molar percent enrichment (MPE) of ^13^C for the indicated metabolites. (F and G) Molar percent enrichment (MPE) of ^13^C for the indicated metabolites over a time course of 16.8 mM U-^13^C glucose tracing in juvenile islets. Glc, glucose; n=4-7 per group; error bars represent ± S.E.M.; *p < 0.05, **p < 0.01, ***p < 0.001 by unpaired two tailed t-test. See also Figure S4 and Table S7.

To validate this method, we determined the changes in labeling and abundance of central carbon metabolites between 2.8 mM and 16.8 mM glucose concentrations. This approach enabled detection of many TCA cycle metabolites, several amino acids, and pyruvate (**Figures 4B, 4C, S4A and S4B**). Glucose stimulation caused an expected increase in the abundance of pyruvate, citrate, fumarate, and malate, while amino acid pools were more stable (**Figures 4B, 4C, S4A and S4B**), as reported in β -cell lines (Mugabo et al., 2017). The metabolic fate of glucose was determined by the rate of ^13^C incorporation into the aforementioned metabolites. Pyruvate and TCA metabolites were labeled with ^13^C, indicating that glucose directly contributes to the accumulation of these metabolites during glucose stimulation (**Figures 4D and S4C**). Amino acids derived from glycolysis (glycine, alanine) and the TCA cycle (glutamate, aspartate) were also labeled (**Figures 4E and S4D**).

Over time, isotope labeling plateaus which is considered steady state. Prior to steady state, metabolite labeling and/or abundance are still changing, allowing for detection of differences in the rate of labeled metabolite accumulation. To determine if one hour of U-^13^C glucose labeling corresponds to steady state or dynamic tracer labeling, we performed time course studies of islets incubated with 16.8 mM U-^13^C glucose. At 60 minutes, the aforementioned metabolites had not reached isotopic steady state U-^13^C labeling (**Figures 4F and 4G**, p = 0.07 for fumarate, Student‘s t-test between 60 and 90 minutes), with the exception of pyruvate (p = 0.8, Student’s t-test between 60 and 90 minutes), indicating that a one-hour trace corresponds to the kinetic labeling phase for these metabolites. This labeling dynamic is in agreement with published tracing data from insulinoma cells, indicating rapid labeling of glycolytic metabolites (e.g. pyruvate) followed by progressive labeling of TCA metabolites for up to several hours (Alves et al., 2015). Considering both metabolite pool sizes and degrees of isotopic labeling, this data reveals pronounced increases in contributions of glucose-derived carbon to citrate and malate pools during glucose stimulation (**Figures 4B, 4D, S4A and S4C**). Isotopologue distributions reflect the extent of ^13^C incorporation on a per-molecule basis. During a U-^13^C glucose trace, labeling of TCA metabolites can occur via combinations of reactions acting upon pyruvate (e.g. pyruvate dehydrogenase and pyruvate carboxylase) and through multiple turns of the TCA cycle (Alves et al., 2015). Glucose stimulation increased the extent of ^13^C incorporation into citrate and malate, giving rise to fully-labeled isotopologues for each of these metabolites (**Figures 5A-D, S5A and S5B**). The accumulation of extensively labeled malate as well as citrate (**Figures 4B, 4D, 4F, S4A, S4C and 5A-D**) supports prior studies, which have indicated rapid anaplerosis of glucose-derived carbon in β-cells (Farfari et al., 2000; Flamez et al., 2002). Thus, our experimental design allows for glucose tracing and targeted metabolomics in islets, enabling the assessment of glucose metabolism in islet endocrine cells in their native context.

**Figure 5.**
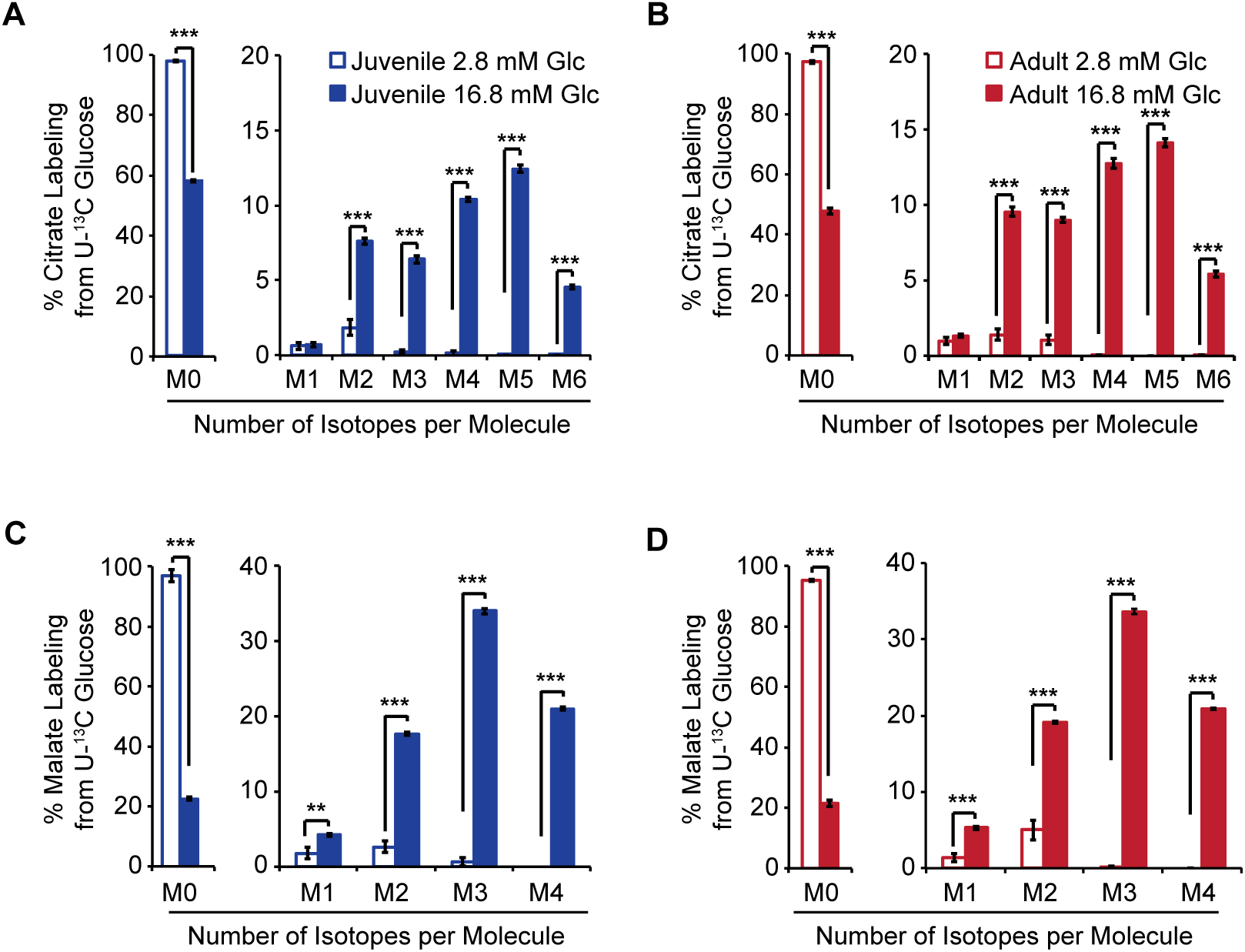
Isotopologue distributions following U-^13^C glucose tracing and targeted metabolomics in islets from juvenile and adult mice. (A-D) Isotopologue distributions of ^13^C for citrate (A and B) and malate (C and D) in juvenile (A and C) and adult (B and D) islets following a one-hour trace with 2.8 mM or 16.8 mM U-^13^C glucose. Isotopologue distributions are expressed as % of the total metabolite pool containing the indicated number of labeled atoms. Glc, glucose; n=4-6 per group; error bars represent S.E.M.; *p < 0.05, **p < 0.01, ***p < 0.001 by unpaired two tailed t-test. See also Figure S5 and Table S7.

### Enhanced entry of glucose-derived carbon into the TCA cycle and accumulation of TCA metabolites in response to glucose in adult islets

We next sought to investigate whether juvenile and adult islets exhibit differences in glucose metabolism that could account for the increase in GSIS with age. Specifically, we explored the hypothesis that quantitative changes to glucose metabolism at the level of the TCA cycle could contribute to the enhancement in glucose-mediated activation of the amplifying pathway in adult islets (**Figure 3A**). Supporting the idea that reactions leading to the production of metabolic coupling factors could be stimulated with age, we observed higher abundance of mitochondrial TCA enzymes (Pdha1, Pdhb, Dlat, Cs, Idh3a, Idh3b, Idh3g, Dhtkd1, Dld, Sdha, and Fh1), the anaplerotic enzyme Pcx, and the pyruvate cycling enzymes Mdh1, Me1, and Acly in islets from 1-year-old compared to 4-week-old mice (**Figures 6A and 6B**). Among the transporters that regulate GSIS by shuttling metabolites between mitochondria and the cytosol (Guay et al., 2007; Huypens et al., 2011; Odegaard et al., 2010), the citrate/isocitrate carrier (CIC) Slc25a1 was expressed at higher levels in adult islets (**Figure 6A**). To assess whether these proteins are transcriptionally regulated in β-cells with age, we analyzed published RNA-seq data of FACS-sorted β-cells isolated at similar time points (Avrahami et al., 2015). β-cells exhibited age-associated increases in mRNA levels of some (i.e. *Acly, Mdh1*, and *Sdha*), but not all, of these genes (**Figure S6A**).

**Figure 6.**
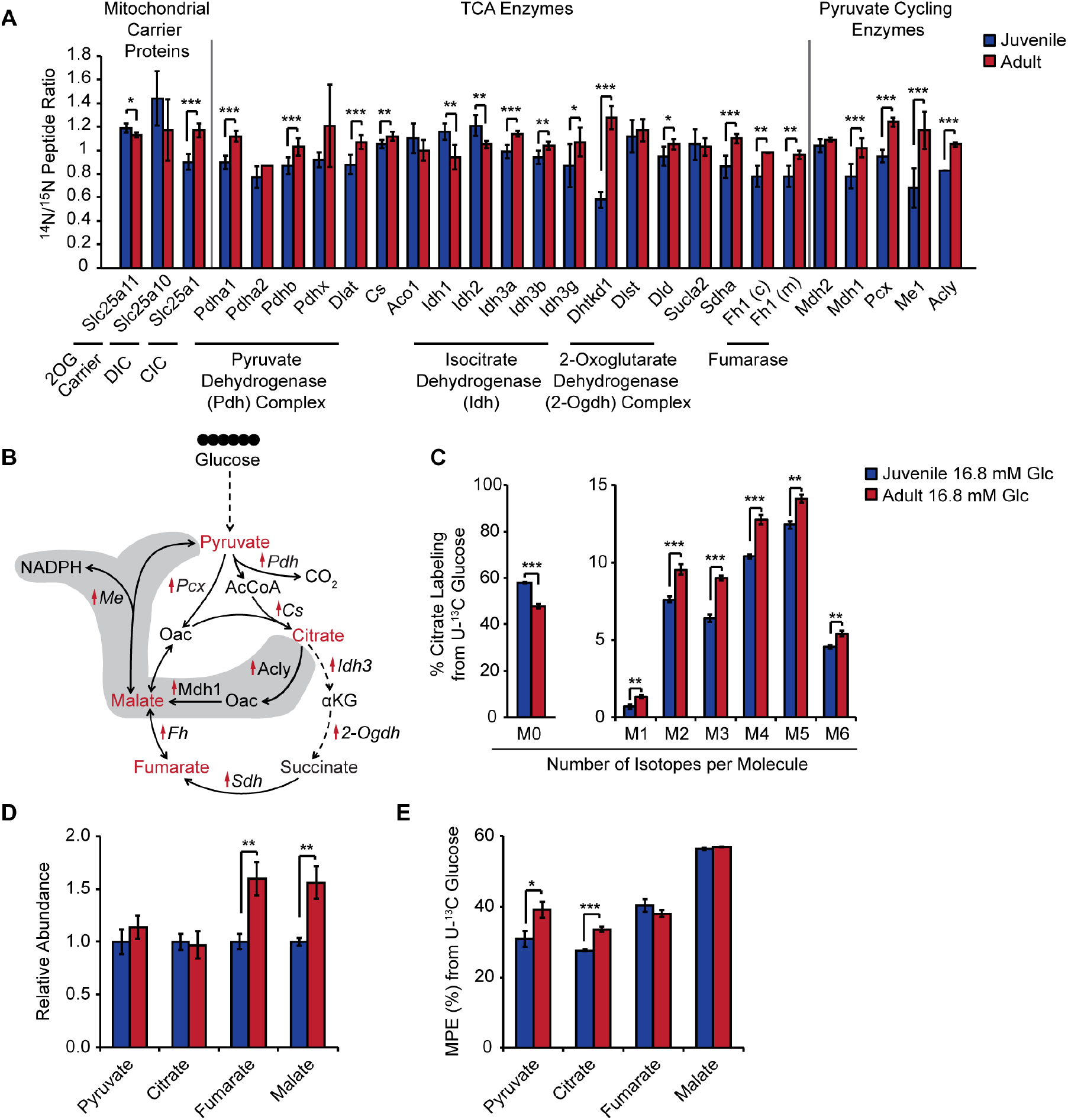
Increased abundance of mitochondrial metabolic enzymes associates with accelerated glucose metabolism in adult islets. (A) Relative protein abundances for the indicated enzymes and transporters in juvenile and adult islets. DIC, dicarboxylate carrier; CIC, citrate/isocitrate carrier. (B) Schematic for TCA and pyruvate cycling enzymes upregulated in adult islets. Upregulated proteins are indicated with red arrows. Intermediary metabolites detected (A) by GC-MS in islets are shown in red. Key reactions of the pyruvate-citrate and pyruvate-malate cycles are shaded. (C) Isotopologue distribution of ^13^C for citrate following a one-hour trace with 16.8 mM U-^13^C glucose in juvenile and adult islets. (D) Abundances of the indicated metabolites, expressed relative to metabolite levels of juvenile islets. (E) Molar percent enrichment (MPE) of ^13^C for the indicated metabolites. Glc, glucose; n=4-6 per group; error bars represent ± S.E.M.; *p < 0.05, **p < 0.01, ***p < 0.001 by unpaired two tailed t-test. See also Figure S6 and Table S7.

To determine whether these changes in abundance of TCA enzymes, pyruvate cycling enzymes, and mitochondrial transporters coincide with changes in glucose metabolism, we compared metabolite abundances and ^13^C labeling in juvenile and adult islets during stimulation with 16.8 mM U-^13^C glucose (**Figures 6C-E, S6B, and S6C**). Isotopologue distributions of citrate revealed higher ^13^C accumulation for all citrate isotopologues during glucose stimulation in adult islets (**Figure 6C**), reflecting increased entry of glucose-derived carbon into the TCA cycle. Furthermore, adult islets accumulated TCA metabolites fumarate and malate to a greater extent than juvenile islets (**Figure 6D**). These changes in TCA metabolites could be driven in part by higher abundance of metabolic enzymes, including pyruvate metabolic enzymes (Pdha1, Pdhb, and Pcx) and TCA enzymes downstream of citrate (Sdha, Fh1, and components of the 2-Ogdh complex). Together, these observations show that age-related increases in mitochondrial metabolic enzymes are associated with enhanced accumulation of TCA metabolites in response to glucose.

### Increased production of glucose-dependent metabolic coupling factors in adult islets

We predicted that enhanced accumulation of TCA metabolites in adult islets could increase production of metabolic coupling factors in the cytosol to augment GSIS. The accumulation of TCA metabolites during glucose stimulation has been attributed to a high rate of anaplerosis driven by the Pcx reaction (Farfari et al., 2000; Flamez et al., 2002). Of TCA metabolites, malate in particular has been implicated in the regulation of GSIS, based on evidence that treatment of islets with a cell-permeable malate analog augments GSIS (Lu et al., 2002) and that the dicarboxylate carrier (DIC), which exports malate from the mitochondria, is required for maximal insulin secretion (Huypens et al., 2011). Mechanistically, malate is a shared metabolite for two well-established pathways of coupling factor generation: pyruvate-malate cycling and pyruvate-citrate cycling. Both cycles involve conversion of pyruvate to oxaloacetate with the Pcx reaction and then the cycles diverge based on the route of pyruvate regeneration. The pyruvate-malate cycle proceeds with reductive metabolism of Pcx-derived oxaloacetate into malate, which is exported into the cytosol then acted upon by Me1 to produce the coupling factor NADPH as pyruvate is regenerated (**Figure 6B**) (MacDonald, 1995). The pyruvate-citrate cycle involves export of citrate into the cytosol, hydrolysis of citrate into oxaloacetate and acetyl-CoA, and conversion of oxaloacetate into malate. Malate can then either serve as a substrate for Me1 or be imported into the mitochondria to re-enter the TCA cycle, though the extent to which malate follows these two paths has been debated (Alves et al., 2015; Prentki et al., 2013). The pyruvate-citrate cycle can produce the coupling factor NADPH (via Me1) as well as cytosolic acetyl-CoA (via Acly), which can then be converted to the coupling factor malonyl-CoA. Since we observed increased abundance of enzymes specific to the pyruvate-citrate cycle (Acly and Mdh1) as well as Me1 common to both cycles (**Figures 6A and 6B**), we predicted that increased pyruvate cycling could contribute to age-dependent enhancement of the amplifying pathway of insulin secretion. To test this possibility, we first measured enzymatic activities of Pcx and Me1, which are two key enzymes shared between the pathways, in islet lysates. Indeed, activities of both enzymes were higher in islets from adult mice (**Figures 7A and 7B**), suggesting that age-dependent increases in protein abundance of these enzymes result in greater enzymatic activities. Both of the aforementioned pyruvate cycles contribute to GSIS via production of the coupling factor NADPH in the Me1 reaction (**Figures 6B and 7C**). NADPH has been shown to directly stimulate GSIS when applied intracellularly (Ivarsson et al., 2005). Because Me1 activity is increased and its substrate malate is more abundant in adult islets, we predicted that glucose-stimulated NADPH generation would be higher in islets from adult mice. Measurement of NADPH levels during glucose stimulation indeed revealed increased NADPH content and a higher NADPH/NADP+ ratio in islets from adult than from juvenile mice (**Figure 7D**). Together, these results indicate increased entry of glucose-derived carbon into the TCA cycle and enhanced production of the metabolic coupling factor NADPH in adult islets. The resulting increase of NADPH could contribute to age-dependent enhancement of GSIS.

**Figure 7.**
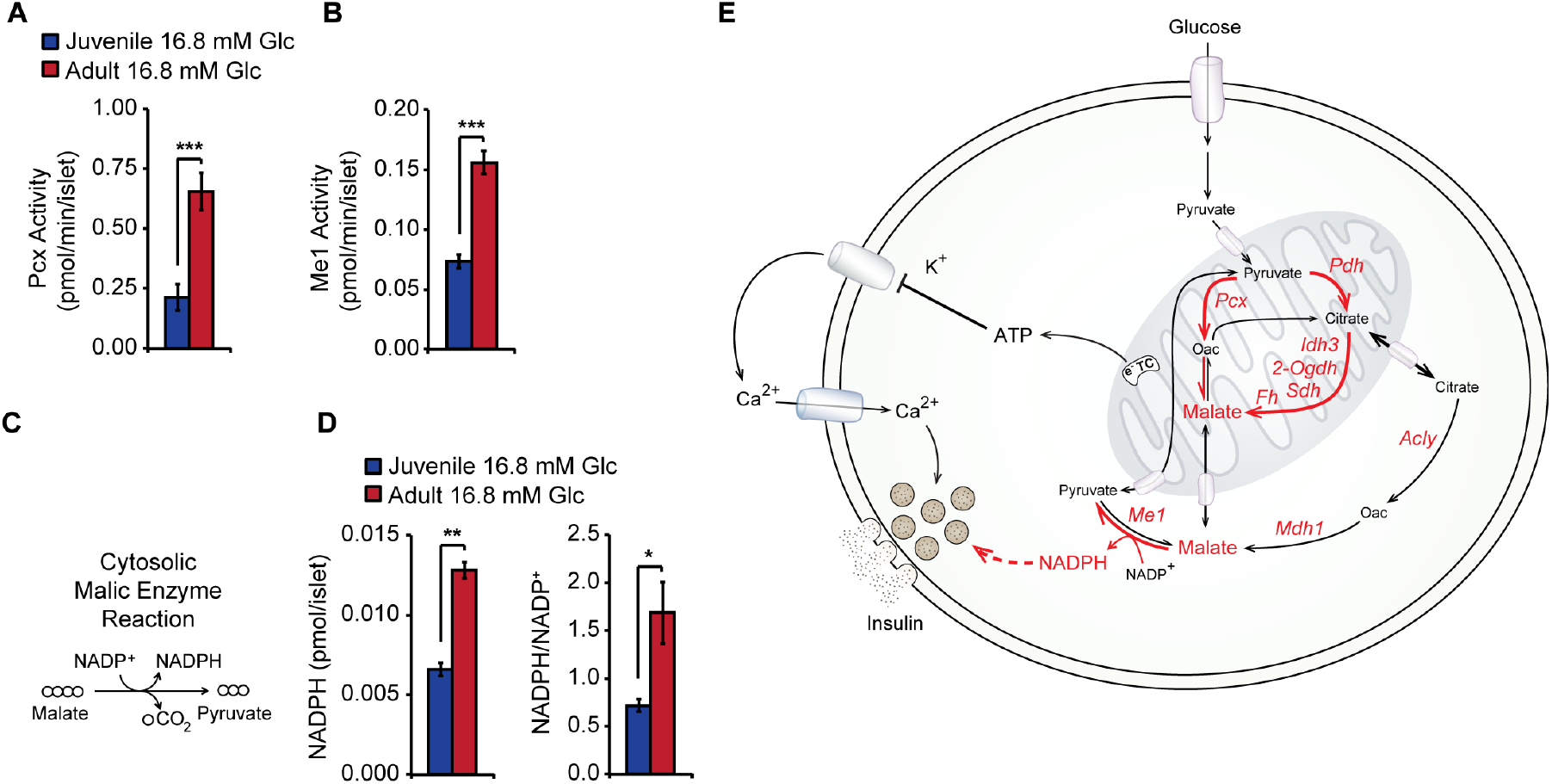
Higher activities of pyruvate cycling enzymes and increased NADPH production in adult islets. (A) Pyruvate carboxylase (Pcx) activity after a one-hour incubation in 16.8 mM glucose in lysates of juvenile and adult islets. (B) Malic enzyme 1 (Me1) activity after a one-hour incubation in 16.8 mM glucose in lysates of juvenile and adult islets. (C) Schematic of the cytosolic malic enzyme reaction as measured in (B). (D) NADPH levels and NADPH/NADP+ ratio after a one-hour incubation in 16.8 mM glucose in juvenile and adult islets. (E) Summary of age-dependent changes in glucose metabolism and coupling factor generation. Red arrows and text indicate reactions, enzymes, and metabolites that increase in abundance and/or activity with age in islets as inferred by nutrient tracing, quantitative proteomics, targeted metabolomics, and measurement of enzymatic activity. Dashed lines indicate coupling factor activation of the amplifying pathway of insulin secretion. Glc, glucose; n=4-11 per group; error bars represent ± S.E.M.; *p < 0.05, **p < 0.01, ***p < 0.001 by unpaired two tailed t-test; n.s., not significant.

## Discussion

It has been increasingly recognized that β-cell regeneration and function exhibit age-related changes (Arda et al., 2016; Avrahami et al., 2015; Gregg et al., 2016; Helman et al., 2016). Here we show that there is age-dependent regulation at the level of islet protein abundance independent of transcription, in particular of proteins involved in the regulation of cellular fuel metabolism. Using metabolic tracing of glucose, we demonstrate that these changes in the abundance of metabolic enzymes are associated with increased production of intermediate metabolites that regulate GSIS. These intermediate metabolites have been implicated in regulation of the amplifying pathway of insulin secretion, which we show to be more active in islets from adult mice. Prior studies have described age-related changes to the triggering pathway of GSIS at the level of KATP channel conductance (Gregg et al., 2016) and Ca^2+^ influx (Avrahami et al., 2015). Given that the metabolic component of the amplifying pathway accounts for roughly half of GSIS (Henquin, 2009), the here-described age-related changes in islet metabolism are likely a significant contributor to altered insulin secretion with age.

The initiating stimulus for GSIS is ATP produced from glucose metabolism and oxidative phosphorylation (OxPhos). It has been previously suggested that the age-related increase in GSIS is driven by an increase in islet OxPhos (Helman et al., 2016). This conclusion was based on the finding that transgenic expression of the senescence protein p16, which increases in β -cells with age, leads to increased islet oxygen consumption during glucose stimulation. However, we found that glucose oxidation, glucose-stimulated respiration, and maximal respiratory capacity were unchanged in adult islets, implying that the age-related increase in GSIS is not driven by changes in OxPhos. Since the study by Helman and Dor examined oxygen consumption in the context of p16 misexpression but not age progression as such, it would indicate that the p16 misexpression model only partially reflects changes in the islet between juvenile and adult mice.

Our observation that OxPhos is unchanged between juvenile and adult mice suggests that alternative mechanisms promote the age-dependent increase of GSIS. Our targeted metabolomics and glucose tracing approach revealed that production of GSIS-relevant coupling factors, most notably malate and NADPH, is enhanced in adult islets. These coupling factors are known to potentiate GSIS (Ivarsson et al., 2005; Lu et al., 2002) and their increased abundance could therefore explain the observed hyperactivation of the amplifying pathway with age. Consistent with our finding that malate production increases with age, our study provides evidence for increased entry of glucose-derived carbon into the TCA cycle associated with increased abundance of mitochondrial metabolic enzymes and enhanced activity of the anaplerotic enzyme Pcx. Furthermore, we observed increased protein levels and activities of enzymes that directly generate coupling factors, suggesting that increased enzyme abundance as well as substrate availability could accelerate cognate reactions. Together, these observations support a view wherein upregulation of mitochondrial enzymes in adult islets promotes anaplerosis and accumulation of TCA metabolites, which are acted upon by coupling factor-producing enzymes that are also more abundant with age (**Figure 7E**). However, it is important to note the limitations of our tracing approach. The complexity of working with primary islets limits the ability to resolve molar fluxes through specific enzymatic reactions within β -cells, raising the possibility that metabolite accumulation results from reduced consumption. Having observed nearly-identical rates of OxPhos in juvenile and adult islets, changes in TCA metabolite consumption seems unlikely. Nevertheless, metabolic flux analysis of pure β-cell preparations at different ages would be required to definitively attribute the observed differences in metabolite accumulation to changes in individual reaction rates.

The major source of NADPH during glucose stimulation remains controversial (Prentki et al., 2013). NADPH can be produced by cytosolic Me1 (MacDonald, 1995), cytosolic isocitrate dehydrogenase (Idh1) (Ronnebaum et al., 2006), or by the glucose-6-phosphate dehydrogenase (G6pd) reaction of the pentose phosphate pathway. However, since the pentose phosphate pathway is not activated in response to glucose (Schuit et al., 1997), it is unlikely to have a major role in the regulation of GSIS. Furthermore, our proteomics data indicate that Idh1 is reduced with age, while Me1 is increased. Therefore, our results are most consistent with Me1 having a role in age-dependent regulation of NADPH generation in islets. Some studies have argued that Me1 is not expressed or has very low activity in mouse islets (MacDonald, 2002; MacDonald et al., 2009), and Me1 deficiency or knockdown was reported to have no effect on GSIS (Ronnebaum et al., 2008). In contrast, and consistent with other reports (Li et al., 2008; Xu et al., 2008), we clearly detected Me1 in islets by MS, RNA-seq, and enzyme activity measurements. Our data further suggest that the role of Me1 is age-dependent, raising the possibility that studies performed in younger animals could overlook the significance of this enzyme. Genetic approaches to manipulate metabolic enzymes specifically in the context of age progression would be necessary to determine the main route for NADPH generation in older β-cells.

The age-dependent changes described herein could comprise both maturation-related and aging-related processes. Rodent studies have indicated that β-cell functional changes are acquired progressively, beginning at birth and extending through at least several weeks after weaning (Bliss and Sharp, 1992; Blum et al., 2012; Yoshihara et al., 2016). Prior studies in mice have suggested that the age-dependent increase of GSIS plateaus at 6 months of age (Helman et al., 2016). By definition, developmental maturation is completed at sexual maturity (at 6-7 weeks of age in mice) and subsequent changes are to be considered part of the aging process. Therefore, changes between 4 weeks and 1 year of age, such as those identified in the current study and in the one by (Avrahami et al., 2015), may predominantly impact GSIS in the final stages of maturation (between 4-6 weeks of age) or during early aging (6 weeks to 6 months of age). Importantly, similar to our and Avrahami’s (Avrahami et al., 2015) observations in mice, an age-dependent increase of GSIS has also been described when comparing human islets from pre-pubescent and middle-aged donors (Arda et al., 2016). Further investigation is needed to determine precisely when these functional and molecular changes take place.

Prior studies have indicated that β-cells undergo metabolic rewiring at the early stages of functional maturation. Perinatal β-cells exhibit high basal insulin secretion and a very low stimulation index in response to glucose, which gradually increases with age (Blum et al., 2012). These early functional changes are associated with increased glucose oxidation and reduced aerobic glycolysis (Boschero et al., 1990). Previous studies have described changes in mRNA levels of metabolic enzymes that occur concordantly with β-cell functional maturation (Jermendy et al., 2011; Yoshihara et al., 2016), suggesting that transcriptional changes contribute to metabolic rewiring. Our study revealed significant age-associated changes in islet protein abundance of TCA cycle and pyruvate cycling enzymes that are not reflected at the transcript level. Thus, while some of the changes in metabolic enzyme protein levels may be driven by transcriptional regulation, our findings indicate a significant role for posttranscriptional mechanisms in islet metabolic rewiring.

With 6,113 proteins faithfully quantified after in vivo labeling, this study represents the most comprehensive proteomics datasets of islets in their native context. The data provide a rich resource for understanding the effects of age on islet physiology. In addition to metabolic enzymes, we also observed age-related posttranscriptional regulation of proteins involved in RNA splicing and cell cycle control. There is growing evidence that RNA splicing regulates β-cell function (Juan-Mateu et al., 2017) and β-cell proliferation is known to decline with age (Teta et al., 2005). In the future, these proteomics datasets will facilitate discovery of additional mechanisms of β-cell functional regulation and β -cell proliferative decline with age. Such knowledge will help identify novel targets for the treatment of diabetes.

## Acknowledgements

The authors would like to thank A. Carrano for critical reading and feedback, J. Fleischman for technical assistance, and N. Rosenblatt for mouse husbandry. We acknowledge support of the UCSD IGM Genomic Center (P30 DK064391) for RNA-seq, as well as the UCSD TEM Core Facility. This work was supported by National Institutes of Health (NIH) training grants T32GM008666-15 (J.R.B.), F32AG039127 (J.N.S.), and T32DK007494-30 (M. Wortham), Juvenile Diabetes Research Foundation (JDRF) postdoctoral fellowship 3-PDF-2014-193-A-N (M. Wortham), John G. Davies Endowed Fellowship in Pancreatic Research (M. Wortham), NIH grants CA188652 (C.M.M.) and DK068471 (M.S.), and JDRF grant 17-2012-35 (M.S.).

## Author Contributions

J.R.B., M. Wortham, and M.S. conceived the project, analyzed data, and wrote the manuscript. J.R.B. performed islet and sample preparation for MS, RNA-seq, and TEM. J.R.B. performed insulin secretion and mitochondria experiments. M. Wortham, M. Wallace, and C.M. performed metabolite measurements and glucose tracing experiments. J.N.S. and J.R.Y.III performed mass spectrometry, V.A., M. Wortham, and E.S. glucose oxidation experiments, J.R.B., M. Wortham, A.D., and A.M. mitochondrial respirometry experiments, and F.M. and Y.S. bioinformatic analysis. B.L.T. and F.L. contributed to islet experiments. All authors reviewed the manuscript.

## Contact for Reagent and Resource Sharing

Further information and requests for resources and reagents should be directed to and will be fulfilled by the Lead Contact, Maike Sander

### Experimental Model and Subject Details

#### Primary mouse islets

Mouse islets were isolated from cohorts of equal numbers of male and female C57BL/^6^N mice (Charles River). Mice were housed 5 animals per cage on woodchip bedding. For the SILAM cohort, mice were obtained at the age of weaning (3-4 weeks of age) and fed a ^15^N-rich, Spirulina-based diet (Cambridge Isotope Laboratories) for 10-11 weeks. Non-labeled mice were maintained on a standard chow diet (PicoLab Rodent Diet 20 pellets, #5053). The juvenile mouse cohort was 4-weeks-old and the adult cohort was 10-15 months of age. All animal experiments were approved by the Institutional Animal Care and Use Committees of the University of California, San Diego.

### Method Details

#### Mice and Tissue Preparation

For the SILAM cohort, 56 mixed 1:1 male and female mice were weaned at 3 weeks of age and fed a ^15^N-rich, Spirulina-based diet for 10-11 weeks. Islets were isolated and pooled from these SILAM mice using a standard islet isolation protocol as described (Schaffer et al., 2011). Islets were also isolated from 62 4-week-old and 38 1-year-old mixed 1:1 male and female ^14^N non-labeled mice and pooled to produce 4 biological replicate samples per age group. Briefly, Liberase TL (Roche) was perfused into pancreata at a working concentration of 0.655 units/mL through the common hepatic bile duct. Pancreata were then removed and dissociated at 37°C for roughly 15 minutes (dissociation time depends on age and size of pancreas). Islets were separated onto a gradient composed of HBSS (Cellgro) and Histopaque (Sigma) layers. Purified islets were then handpicked under a dissection microscope to minimize acinar cell contamination. Islets were lysed in a buffer containing 10 mM Tris-HCl pH8, 10 mM NaCl, 3 mM MgCl2, 1% NP-40, 1% SDS, 0.5% Sodium Deoxycholate with 1x protease inhibitor cocktail (Roche) and 1 mM PMSF. 40 µg of ^15^N protein was mixed with 40 µg of ^14^N protein for 4 juvenile biological replicates and 3 adult biological replicate samples. A fourth adult biological replicate sample was composed of 20 µg of ^15^N and 20 µg of ^14^N protein.

#### MudPIT LTQ Velos Orbitrap Mass Spectrometry

##### Sample Preparation

Urea (8 M) was added to 80 µg of protein lysates for LCLC-MS/ MS analysis, and extracts were processed with ProteasMAX (Promega) following the manufacturer’s instructions. The samples subsequently were reduced by 5 mM Tris(2-carboxyethyl)phosphine at room temperature for 20 minutes, alkylated in the dark by 10 mM iodoacetamide for 20 minutes, and digested with Sequencing Grade Modified Trypsin (Promega) overnight at 37 °C, and the reaction was stopped by acidification.

##### MudPIT and LTQ Velos Orbitrap MS

The protein digest was pressure-loaded into a 250-µm i.d capillary packed with 2.5 cm of 10-µm Jupiter C18 resin (Phenomenex) followed by an additional 2.5 cm of 5-µm Partisphere strong cation exchanger (Whatman). The column was washed with buffer containing 95% (vol/vol) water, 5% (vol/vol) acetonitrile, and 0.1% formic acid. After washing, a 100-µm i.d capillary with a 5-µm pulled tip packed with 15 cm of 4-µm Jupiter C18 resin was attached to the filter union, and the entire split column (desalting column–union–analytical column) was placed inline with an Agilent 1200 quaternary HPLC and analyzed using a modified 11- step separation described previously (Link et al., 1999; Washburn et al., 2001). The buffer solutions used were 5% (vol/vol) acetonitrile/0.1% formic acid (buffer A), 80% (vol/vol) acetonitrile/0.1% formic acid (buffer B), and 500 mM ammonium acetate/5% (vol/vol) acetonitrile/0.1% formic acid (buffer C). Step 1 consisted of a 90-min gradient from 0–100% (vol/vol) buffer B. Steps 2–11 had a similar profile with the following changes: 5 min in 100% (vol/vol) buffer A, 3 min in X% (vol/vol) buffer C, a 10-min gradient from 0 to 15% (vol/vol) buffer B, and a 108-min gradient from 15–100% (vol/vol) buffer B. The 3-min buffer C percentages (X) were 10, 20, 30, 40, 50, 60, 70, 80, 90, and 100% (vol/vol), respectively, for the 11-step analysis. As peptides eluted from the microcapillary column, they were electrosprayed directly into an LTQ Velos Orbitrap mass spectrometer (Thermo Finnigan) with the application of a distal 2.4-kV spray voltage. A cycle of one full-scan mass spectrum (400–1,800 m/z) at a resolution of 60,000 followed by 15 data-dependent MS/MS spectra at a 35% normalized collision energy was repeated continuously throughout each step of the multidimensional separation. Maximum ion accumulation times were set to 500 ms for survey MS scans and to 100 ms for MS2 scans. Charge state rejection was set to omit singly charged ion species and ions for which a charge state could not be determined for MS/MS. Minimal signal for fragmentation was set to 1,000. Dynamic exclusion was enabled with a repeat count of 1, duration of 20.00 s, list size of 300, exclusion duration of 30.00 s, and exclusion mass with high/low of 1.5 m/z. Application of mass spectrometer scan functions and HPLC solvent gradients were controlled by the Xcaliber data system.

##### Analysis of Tandem Mass Spectra

Protein identification and quantification and analysis were done with Integrated Proteomics Pipeline - IP2 (Integrated Proteomics Applications, Inc., www.integratedproteomics.com/) using Pro-LuCID, DTASelect2, Census, and QuantCompare. Spectrum raw files were extracted into ms1 and ms2 files from raw files using RawXtract 1.9.9 (http://fields.scripps.edu/downloads.php) (McDonald et al., 2004), and the tandem mass spectra were searched against the European Bioinformatic Institute (IPI) mouse protein database (www.ebi.ac.uk/IPI/IPImouse.html, downloaded on January 1, 2009) (Kersey et al., 2004). To estimate peptide probabilities and FDRs accurately, we used a target/decoy database containing the reversed sequences of all the proteins appended to the target database (Peng et al., 2003). Tandem mass spectra were matched to sequences using the ProLuCID {Xu T, 2006 #323} algorithm with 50 ppm peptide mass tolerance for precursor ions and 400 ppm for fragment ions. ProLuCID searches were done on an Intel Xeon cluster running under the Linux operating system. The search space included all fully and half-tryptic peptide candidates that fell within the mass tolerance window with no miscleavage constraint. Carbamidomethylation (+57.02146 Da) of cysteine was considered as a static modification.

The validity of peptide/spectrum matches (PSMs) was assessed in DTASelect (Cociorva et al., 2007; Tabb et al., 2002) using two SEQUEST-defined parameters (Eng et al., 1994), the cross-correlation score (XCorr), and normalized difference in cross-correlation scores (DeltaCN). The search results were grouped by charge state (+1, +2, +3, and greater than +3) and tryptic status (fully tryptic, half-tryptic, and nontryptic), resulting in 12 distinct subgroups. In each of these subgroups, the distribution of Xcorr, DeltaCN, and DeltaMass values for (a) direct and (b) decoy database PSMs was obtained; then the direct and decoy subsets were separated by discriminant analysis. Full separation of the direct and decoy PSM subsets is not generally possible; therefore, peptide match probabilities were calculated based on a nonparametric fit of the direct and decoy score distributions. A peptide confidence of 0.95 was set as the minimum threshold. The FDR was calculated as the percentage of reverse decoy PSMs among all the PSMs that passed the confidence threshold. Each protein identified was required to have a minimum of two peptides that are at least half-tryptic; however, these peptides had to be an excellent match with an FDR less than 0.001 and at least one excellent peptide match. We also require the precursor m/z to be less than or equal to 10 ppm from the theoretical m/z. After this last filtering step, we estimate that the protein FDRs were below 1% for each sample analysis.

Each dataset was searched twice, once against light and then against heavy protein databases. After the results from SEQUEST were filtered using DTASelect2, ion chromatograms were generated using an updated version of a program previously written in Yates laboratory (MacCoss et al., 2003). This software, called “Census (Park et al., 2006), is available from the authors for individual use and evaluation through an Institutional Software Transfer Agreement (see http://fields.scripps.edu/census for details).

First, the elemental compositions and corresponding isotopic distributions for both the unlabeled and labeled peptides were calculated, and this information was then used to determine the appropriate m/z range from which to extract ion intensities, which included all isotopes with greater than 5% of the calculated isotope cluster base peak abundance. MS1 files were used to generate chromatograms from the m/z range surrounding both the unlabeled and labeled precursor peptides.

Census calculates peptide ion intensity ratios for each pair of extracted ion chromatograms. The heart of the program is a linear least-squares correlation that is used to calculate the ratio (i.e., slope of the line) and closeness of fit [i.e., correlation coefficient (r)] between the data points of the unlabeled and labeled ion chromatograms. Census allows users to filter peptide ratio measurements based on a correlation threshold; the correlation coefficient (values between zero and one) represents the quality of the correlation between the unlabeled and labeled chromatograms and can be used to filter out poor-quality measurements. In this study, only peptide ratios with the coefficient correlation values (r2) greater than 0.5 were used for further analysis. In addition, Census provides an automated method for detecting and removing statistical outliers. In brief, SDs are calculated for all proteins using their respective peptide ratio measurements. The Grubbs test (P < 0.01) then is applied to remove outlier peptides. The outlier algorithm is used only when more than two peptides are found in the same protein, because the algorithm becomes unreliable for a small number of measurements.

Final protein ratios were generated with QuantCompare, which uses Log two-fold change and t-test p-value to identify regulated significant proteins. For a protein to be considered for follow up analysis, it had to qualify for plotting onto our volcano scatter plot (**Figure 1B**). The y axis of these volcano plots is the t-test p-value, which requires each protein to be quantified in at least two of the biological replicates (so we can calculate the variance) for both juvenile and adult mice. To show how many measurements our quantitative measures were drawn from, we have provided peptide counts for each protein listed in Tables S1B and S1C (see the columns headed “Total number of quantitated ^14^N/^15^N pairs).

#### Gene Ontology Analysis

Gene Ontology (GO) analysis was performed using the Panther Classification System (Thomas et al., 2003) or the analysis of the entire identified proteome and using Metascape (Tripathi et al., 2015) for the analysis of the significantly changed proteins with age.

#### RNA-Sequencing

Isolated islets were pooled from two mice each at 4 weeks and 1 year of age for a total of three samples per age that were lysed in RLT Buffer. Total RNA was extracted using the RNeasy Micro Kit (Qiagen) per manufacturer’s instructions. TruSeq stranded mRNA libraries were prepared by the UCSD Institute for Genomic Medicine Genomics Center and sequenced using HiSeq2500 Highoutput Run V4 (Illumina). The single-end 50 base pair reads were mapped to the UCSC mouse transcriptome (mm9) by STAR, allowing for up to 10 mismatches. Only the reads aligned uniquely to one genomic location were retained for subsequent analysis. Expression levels of all genes were estimated by Cufflinks (Trapnell et al., 2010) using only the reads with exact matches.

#### Comparative GSEA of Protein and RNA Data

In order to compare differential regulation observed in RNA and proteins, fold changes (FC) representing differences of adult compared to juvenile islet samples were considered for each gene g_i_, with i ranging from one to the number of genes with encoded proteins found in the proteomic dataset. The two sets of fold changes available SF_P_={FC_i_^P^} and SF_R_={FC_i_^R^}, accounting for expression changes at the level of proteins and RNA, respectively, were subsequently scored for global concordance by computing their Spearman rank correlation. A second comparison of the two data sets focused on gene sets, i.e. groups of genes participating in known biological pathways. A similar approach to the one described above was adopted to collect measures of the differential changes in all genes found in both datasets, this time by taking into account p-values and performing a separate analysis for up- and downregulated genes.

Considering only genes upregulated in adult compared to juvenile islets, two quantitative signatures SP_P_-UP={p_k_^P^} and SP_R_-UP={p_k_^R^} were obtained, with p_k_ representing the significance (p-value) of the k-th upregulated gene and the two sets SP_P_-UP and SP_R_-UP accounting for changes at the level of proteins and RNA, respectively.

Genes in the signatures were first mapped to Human Ensembl gene identifiers using Biomart (Kasprzyk, 2011) and subsequently analyzed with GSEA (Subramanian et al., 2005). In details, the first step of GSEA analysis allowed ordering all up-regulated genes according to the differences of their p-values in SP_P_-UP and SP_P_-UP, thus obtaining a ranked list *L* with the top-ranked elements being genes for which a more significant change (p-value) in proteins, compared to RNA, was observed.

Finally, for each canonical pathway from collection C2 of the MSigDB database (Liberzon et al., 2011), the Enrichment Score (ES) and correspondent enrichment p-values were computed with GSEA to identify pathways for which gene members were found at the top (positive ES) or at the bottom (negative ES) of L. Pathways (FDR < 0.25) enriched for up-regulated genes with more significant changes in proteins (positive ES) or in RNA (negative ES), were considered for further analysis and interpretation. Analogous signatures SP_P_-DOWN and SP_R_-DOWN were derived for down-regulated genes and the approach described above was used to compare their changes in proteins and RNA.

#### Comparative Network-Based Analysis of Protein and RNA Data

Genes corresponding to regulated proteins with p<0.05 and abs(FC)≥1.2 (n=872) were connected to their top three interactors through the protein interaction repository STRING (version 10.0, http://string-db.org/). Only protein associations with combined confidence score > 0.4 were retained and used to assign weights to each network link. Functional categories related to the entire set of proteins included in the network and links between each pair of categories were identified with Metascape as described in http://metascape.org/. In details, all the statistically enriched terms identified from GO, KEGG and hallmark gene sets were hierarchically clustered into a tree based on Kappa-statistical similarities among their gene memberships. A 0.3 kappa score was applied as the threshold to define clusters, each of them representing a group of similar functional categories. A subset of representative terms from each cluster was automatically selected by Metascape and converted into a network, where terms with similarity > 0.3 are connected by edges. More specifically, terms with the best p-values from each of the clusters were depicted as network nodes, with the constraint of having a maximum of 15 terms per cluster and 250 terms in total.

The same network building procedure was adopted for significant genes with p<0.05 and abs(FC)≥1.2 for which a correspondent entry was found in the protein data (n=331). Then, the two networks of protein-related and RNA-related functional categories were compared by adopting a protein-centered analysis. First, nodes belonging to both protein and RNA networks were identified and scored for differences in enrichment p-values. For the *j-th* node representing a functional category enriched enriched in both protein and RNA networks, a score *S_j_* was computed as *S_j_ = log(p_j_^P^) – log(p_j_^R^)*, with *p_j_^P^* and *p_j_^R^* representing the enrichment p-values assigned by Metascape to the j-th category in the protein and in the RNA data, respectively. Node sizes were adjusted proportionally to *S*, with larger nodes representing a more significant enrichment at the protein level. The same procedure was used for all remaining nodes, i.e. categories enriched only at the protein level. Scores were computed in this case by setting *p_j_^R^* = 1, thus obtaining node sizes proportional to the significance of these categories in the protein data.

#### Glucose Stimulated Insulin Secretion

Glucose stimulated insulin secretion (GSIS) assays were performed as previously described (Schaffer et al., 2011). Briefly, islets were isolated and pooled from seven to eighteen mice per group, then incubated overnight in RPMI 1640 supplemented with 8 mM glucose, 10% FBS, 2 mM L-glutamine, 100 U/mL Pen/Strep, 1 mM sodium pyruvate, 10 mM HEPES, and 0.25 mg/mL amphoterecin B. The next day, islets were washed and pre-incubated for 1 hour in Krebs-Ringers-Bicarbonate-HEPES (KRBH) buffer (130 mM NaCl, 5 mM KCl, 1.2 mM CaCl_2_, 1.2 mM MgCl_2_, 1.2 mM KH_2_PO_4_, 20 mM HEPES pH 7.4, 25 mM NaHCO_3_, and 0.1% bovine serum albumin) supplemented with 2.8 mM glucose solution at 37°C with 5% CO_2_. Afterward, groups of 10 islets that were size-matched between groups were transferred to a 96 well plate into KRBH solutions containing 2.8 mM glucose, 16.8 mM glucose, 10 mM leucine/glutamine with 16.8 mM glucose, or 75 µM palmitate with 16.8 mM glucose. Palmitate was non-covalently conjugated to fatty acid-free BSA at a 6:1 molar ratio and was added to KRBH buffer at a total concentration of 75 µM. To clamp the triggering pathway and measure the amplifying pathway, islets were incubated in 100 µM diazoxide and 30 mM KCl and 2.8 mM glucose, 16.8 mM glucose, 10 mM leucine/glutamine with 2.8 mM glucose, or 75 µM palmitate with 2.8 mM glucose. After incubation for 1 hour at 37°C in 5% CO_2_, supernatant was collected and islets were lysed overnight in a 20% acid:80% ethanol solution. Insulin was then measured in supernatants and lysates using a mouse insulin ELISA (ALPCO). Secreted insulin was calculated as percentage of total insulin content per hour.

#### Electron Microscopy

For electron microscopy sample preparation, a fixative containing 2% paraformaldehyde, 2.5% glutaraldehyde, 3 mM CaCl_2_ prepared in 0.1M sodium cacodylate buffer to a final pH 7.4 was used on isolated islets pooled from 6 mice and stored at 4°C overnight. Islets were postfixed in 1% osmium tetroxide in 0.15 M cacodylate buffer for 1 hour and stained en bloc in 1% uranyl acetate for 1 hour. Samples were dehydrated in ethanol, embedded in Durcupan epoxy resin (Sigma-Aldrich), sectioned at 50 to 60 nm, and picked up on Formvar and carbon-coated copper grids. Sections were then stained with 2% uranyl acetate for 5 min and Sato’s lead stain for 1 min. Grids were viewed using a Tecnai G2 Spirit BioTWIN transmission electron microscope equipped with an Eagle 4k HS digital camera (FEI, Hilsboro, OR). To quantify mitochondria number, 50-60 β-cells were randomly imaged from multiple image planes and multiple sections at a 1900x magnification. Individual β-cells were identified by the observation of insulin granules within defined cell borders visualized by cell-cell junctions. Mitochondria were identified and counted by the observation of cristae structures. And to quantify mitochondrial length, the longest distance between two points was measured using Image J (Rasband, 1997-2016). In 10 randomly selected β-cells, 49-59 mitochondria were measured for length. Quantifications of mitochondria number and length were performed blinded.

#### Mitochondrial DNA Quantification

To measure mitochondrial DNA copy number, total DNA from isolated islets from 4-week-old and 1-year-old mice was isolated using DNeasy Blood & Tissue Kit (Qiagen) according to the manufacturer’s instructions. Mitochondrial DNA (mtDNA) and nuclear DNA (nDNA) content were determined by quantitative real-time PCR using A CFX96 real-time system (BioRad) with specific primers for the mitochondrial cytochrome c oxidase subunit II (*Cox2*) gene and the nuclear gene *Rsp18* using the following primer sequences: mt-*Cox2*: 5′-ATAACCGAGTCGTTCTGCCAAT-3′ and 5′-TTTCAGAGCATTGGCCATAGAA-3′, *Rsp18*: 5′- TGTGTTAGGGGACTGGTGGACA-3′ and 5′- CATCACCCACTTACCCCCAAAA-3′. The ratio of mtDNA to nDNA content was calculated for each time point. Experiments were performed on 3 biological replicate samples.

#### Enzyme Activity Measurements

Islets were isolated and pooled from 8-10 mice per group for each assay, allowed to recover overnight, and incubated in 2.8 mM glucose as for GSIS assays. Following incubation in 2.8 mM glucose, islets were transferred to KRBH supplemented with 16.8 mM glucose and immediately seeded into wells of non-treated 12-well plates (Falcon #351143) at a density of 150 (for pyruvate carboxylase) or 200 (for malic enzyme) size-matched islets per well in a final volume of 500 µl. Islets were incubated for 1 hour at 37°C with 5% CO_2_, then islets were transferred to Eppendorf tubes on ice, centrifuged at 500 x *g* for 2 min at 4°C, washed in 1 mL ice cold DPBS, pelleted again, and resuspended in lysis buffer (10 mM HEPES pH 7.4, 250 mM sucrose, 2.5 mM EDTA, 2mM L-Cysteine, and 0.02% bovine serum albumin). Lysates were prepared by passaging 20x through a 26 gauge needle, incubating on ice for 10 min, then vortexing for 10 s.

Pyruvate carboxylase activity measurements were performed using a protocol adapted from (Liu et al., 1999). Briefly, 16 µL of islet lysates were loaded into a 200 µL reaction mixture containing 134 mM Triethanolamine, 9 mM MgSO_4_, 7 mM sodium pyruvate, 0.23 mM NADH, 2mM ATP, 0.16 mM Acetyl-CoA, 21mM KHCO_3_, 0.12% BSA, and 5U of malate dehydrogenase enzyme. The reaction was carried out at 30°C, and absorbance at 340 nm was monitored for 25 min to determine the decrease in absorbance over time (ΔA_340_), which was subsequently used to calculate the reaction rate.

Malic enzyme activity measurements were performed using a protocol adapted from (Xu et al., 2008). Briefly, 32 µL of islet lysates were loaded into a 216 µL reaction mixture containing 67 mM Triethanolamine, 4 mM MgCl_2_, 1 mM L-Malate, and 0.23 mM NADP^+^. The reaction was carried out at 37°C, and absorbance at 340 nm was monitored for 25 min to determine the increase in absorbance over time (ΔA_340_), which was subsequently used to calculate the reaction rate.

#### NADPH/NADP^+^ Measurements

NADPH and NADP^+^ were quantified by the enzyme cycling method (Sigma MAK038). Briefly, islets were isolated and pooled from 6 mice per group, allowed to recover overnight, and incubated in 2.8 mM glucose as for GSIS assays. Following incubation in 2.8 mM glucose, islets were transferred to KRBH supplemented with 16.8 mM glucose and immediately seeded into wells of non-treated 12-well plates (Falcon #351143) at a density of 200 size-matched islets per well in a final volume of 500 µl. Islets were incubated for 1 hour at 37°C with 5% CO_2_, then islets were transferred to Eppendorf tubes on ice, centrifuged at 500 x *g* for 2 min at 4°C, washed in 1 mL ice cold DPBS, pelleted again, and resuspended in extraction buffer (Sigma MAK038). Lysates were prepared by passaging 20x through a 26 gauge needle, incubating on ice for 10 min, then vortexing for 10 s. Debris was pelleted by centrifugation at 13,000 x g for 10 min at 4°C, then supernatant was harvested for immediate nucleotide measurements. NADP^+^ was decomposed in half of each sample by heating at 60°C for 30 min, then NADPH and NADP_total_ concentrations were determined according to the manufacturer’s instructions. NADP^+^ concentration was determined by subtracting NADPH from NADP_total_.

#### Islet Respirometry

Respirometry was performed using a protocol adapted from (Wikstrom et al., 2012). Briefly, islets were isolated and pooled from 10 mice per group, then allowed to recover overnight as for GSIS assays. Islets were then starved in Seahorse XF DMEM Assay Medium supplemented with 1% FBS, 2.8 mM glucose, and 5 mM HEPES (final pH of 7.4) at 37°C without CO_2_ for 45 min. Following starvation, islets were seeded at a density of 100 size-matched islets per well into the depressions of XF24 Islet Capture Microplates containing Seahorse XF DMEM assay medium supplemented as above, then capture screens were inserted into wells. The plate was then loaded into the Seahorse Bioscience XF24 Extracellular Flux Analyzer heated to 37°C, and oxygen consumption was measured sequentially for the following states: basal (2.8 mM glucose), nutrient-stimulated (16.8 mM glucose or 10 mM leucine/glutamine), ATP synthase-independent (5 µM oligomycin), and non-mitochondrial (0.5 µM rotenone and 2 µM antimycin) respiration. Maximum uncoupled respiration was measured by treating with 2 µM FCCP after the addition of leucine/glutamine and oligomycin. Following respirometry, wells were imaged and relative islet area was calculated using ZEN. Oxygen consumption rates for each well were normalized to islet area relative to that of all wells.

#### U-^13^C Glucose Tracing and Metabolite Measurements

Islets were isolated, pooled from 10 mice per group, and allowed to recover overnight as for GSIS assays. Islets were starved for 1 hour in KRBH supplemented with 2.8 mM glucose as for GSIS assays. Islets were then transferred to petri dishes containing U-^13^C-labeled glucose as indicated and immediately seeded into wells of non-treated 12-well plates at a density of 100 size-matched islets per replicate in a final volume of 500 µl KRBH containing basal (2.8 mM) or stimulatory (16.8 mM) concentrations of labeled glucose. Tracing was performed for the indicated durations at 37°C with 5% CO_2_. Following the trace, islets were rapidly chilled by swirling plate to resuspend islets then transferring entire contents of each well to an Eppendorf tube on ice. Islets were then pelleted by centrifugation at 500 x *g* for 2 min at 4°C, washed in 1 mL of ice cold 0.9% NaCl, centrifuged as before, then pellets were frozen in a dry ice/ethanol bath and stored at −80°C.

Metabolites were extracted from pellets using a bligh and dyer-based methanol/chloroform/water extraction with inclusion of norvaline as a polar internal standard. Briefly, 250 µl MeOH, 250 µl CHCL_3_, 100 µl water containing norvaline were added to pellets. This was vortexed for 10 minutes followed by centrifugation at 10,000 g for 5 minutes at 4°C. The upper phase was separated and dried under vacuum at 4°C until dry. Polar metabolites were derivatized in 2% (w/v) methoxyamine hydrochloride in pyridine and incubated at 45°C for 60 minutes. Samples were then silylated with N-tertbutyldimethylsilyl-N-methyltrifluoroacetamide (MtBSTFA) with 1% tert-butyldimethylchlorosilane (tBDMCS) at 45°C for 30 minutes. Polar derivatives were analyzed by GC-MS using a DB-35MS column (30m x 0.25 mm i.d. x 0.25 μm) installed in an Agilent 7890B gas chromatograph (GC) interfaced with an Agilent 5977A mass spectrometer (MS) with an XTR EI source using the following temperature program: 100 °C initial, increase by 3.5 °C/min to 255 °C, increase by 15 °C/min to 320 °C and hold for 3 min. The % isotopologue distributions of each polar metabolite was determined and corrected for natural abundance using in-house algorithms adapted from (Fernandez et al., 1996). Negative isotopologue distributions resulting from natural abundance corrections were rounded to zero. The metabolite ion fragments used are summarized below:

| Metabolite | Carbons | Formula | m/z |
| --- | --- | --- | --- |
| Pyruvate | 1,2,3 | C6H12O3NSi | 174 |
| Alanine | 1,2,3 | C11H26O2NSi2 | 260 |
| Glycine | 1,2 | C10H24O2NSi2 | 246 |
| Aspartate | 1,2,3,4 | C18H40O4NSi3 | 418 |
| Glutamate | 1,2,3,4,5 | C19H42O4N1Si3 | 432 |
| Citrate | 1,2,3,4,5,6 | C20H39O6Si3 | 459 |
| Malate | 1,2,3,4 | C18H39O5Si3 | 419 |
| Fumarate | 1,2,3,4 | C12H23O4Si2 | 287 |

Molar percent enrichment (MPE) represents the percent of ^13^C-labeled carbon in each metabolite pool and was calculated as follows:

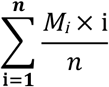

where *n* is the number of carbon atoms in the metabolite and *M_i_* is the relative abundance of the *i*th isotopologue.

#### Glucose Oxidation and Utilization Rate Measurements

Glucose oxidation and utilization rates were determined essentially as described in (Peyot et al., 2010). Briefly, islets were isolated and pooled from 8 mice per group. Islets were allowed to recover overnight, and then starved for 1 hour in KRBH supplemented with 2.8 mM glucose as for GSIS assays. Following starvation, islets were allocated to tubes (50 size-matched islets/tube) and incubated in KRBH containing either 2.8 or 16.8 mM glucose spiked with 1.5 µCi each of U-^14^C glucose and 5-^3^H glucose for 90 min at 37°C. Islets were then treated with 10 µM antimycin-A/rotenone and 5 mM potassium cyanide in citrate/NaOH buffer (400 mM, pH 4.9). ^14^CO_2_ generated during incubation was trapped with KOH and quantified by scintillation counting. ^3^H_2_O generated during incubation was allowed to evaporate and condense with 1M HCl for 40 hours at room temperature and was then quantified by scintillation counting.

## Quantification and Statistical Analysis

For islet experiments, islets were pooled from different mice where n represents the number of biological replicates using islet pools (i.e. 2 mice per 1 biological replicate, n=1). For electron microscopy, n represents the number of individual β-cells from randomly selected images taken from 3 separate sections of a sample of islets pooled from 6 mice. Quantifications of mitochondria number and length were performed blinded using Image J. The exact values of n are reported in the above methods and/or figure legends.

All non-computational analyses were performed with Excel (Microsoft Excel 2011). Data are presented as mean ± standard error of the mean (S.E.M.). Statistical comparisons were completed with unpaired, two-tailed student’s t-test. Significance was defined as p-value < 0.05.

Computational analyses were performed using Bioconductor packages (Gentleman et al., 2004) and other available bioinformatics tools listed below. Proteins were considered significantly regulated with a ratio ≥1.2 and p-value <0.05, estimated with ANOVA and genes were considered significantly regulated with a ratio ≥1.2 and p-value <0.05, estimated with the software Cuffdiff. Concordance of fold changes at the level of proteins and RNA in individual genes was measured with Spearman rank correlation. Groups of genes with more significant coordinated activity at the protein or at the RNA level were identified by running GSEA on significance p-values of genes present in both datasets. Canonical pathways from collection C2 of the MSigDB database (Liberzon et al., 2011) were extracted with FDR < 0.25. Significance for GSEA results was assessed with 1000 permutations and FDR was used to correct for multiple testing. The exact thresholds used for categories and genes selection are specified in the results.

## Data and Software Availability

### Data

The proteomics data have been deposited in Massive and Proteome Xchange under the following IDs:

Massive ID: MSV000081577

ProteomeXchange ID: PXD007899

The proteomics data have been deposited in PINT at the following URL: http://sealion.scripps.edu/pint/?project=bec8ab7dcfa133c1e7e16d47873815ec

The RNA-sequencing data have been deposited in NCBI’s Gene Expression Omnibus (Edgar et al., 2002) under the GEO Series accession number GSE104591 at the following URL: https://www.ncbi.nlm.nih.gov/geo/query/acc.cgi?acc=GSE104591

The targeted metabolomics data have been deposited in NCBI’s Metabolomics Workbench as studies ST000964 and ST000965.

### Software

Integrated Proteomics Pipeline - IP2 Integrated Proteomics Applications, Inc.:

www.integratedproteomics.com/

RawExtract 1.9.9: http://fields.scripps.edu/downloads.php

European Bioinformatic Institute (IPI) mouse protein database:

www.ebi.ac.uk/IPI/IPImouse.html

Census: http://fields.scripps.edu/census

Panther Classification System: http://www.pantherdb.org/

Metascape: http://metascape.org/gp/index.html#/main/step1

Cufflink/Cuffdiff: http://cole-trapnell-lab.github.io/cufflinks/cuffdiff/

Biomart: http://www.biomart.org/

Bioconductor: http://bioconductor.org/

Gene Set Enrichment Analysis (GSEA): http://www.broad.mit.edu/gsea

MSigDB database: http://software.broadinstitute.org/gsea/msigdb

STRING: http://string-db.org/

